## Supplementary material for "A high-resolution analysis of arrestin2 interactions responsible for CCR5 endocytosis": Figure 1-figure supplement 1

**Affiliations:**
[1] Biozentrum, University of Basel, CH-4056 Basel, Switzerland

† present address: Center for Life Sciences, Paul Scherrer Institut, CH-5232 Villigen, Switzerland

#### Tables

**Table S1.** CCR5 phosphopeptide sequences and their affinities towards arrestin2. Underlined serine or threonine residues are phosphorylated.

| Phosphopeptide | Sequence | $K_D$ (arrestin2$^{1\text{-}393}$) [μM][a] |
|---|---|---|
| CCR5pp6 | APERA<u>SS</u>VVY<u>T</u>R<u>ST</u>GEQEI<u>S</u>VGL | 45 ± 6 |
| CCR5pp4 | APERA<u>SS</u>VVYTR<u>S</u>TGEQEI<u>S</u>VGL | 199 ± 10 |
| CCR5pp3 | APERASSVVY<u>T</u>R<u>ST</u>GEQEISVGL | 198 ± 6 |

[a]data from Isaikina, Petrovic et al. (2023).

### Figures

A

```
                                        clathrin motif            AP2 motif
arr1 MHPQPEDPARKES.......................YQDANLVFEEFARHNLKDA 388
arr2 MHPKPKEEPP....HREVPENETPVDTNLIELDTN...DDDIVFEDFARQRLKGM 399
arr3 MHPKPHDHIPLPRPQSAAPETDVPVDTNLIEFDTNYATDDDIVFEDFARLRLKGM 399
arr4 IHPKPSHEAAS..........................SEDIVIEEFTRKGEEES 376
                                                β20
```

B

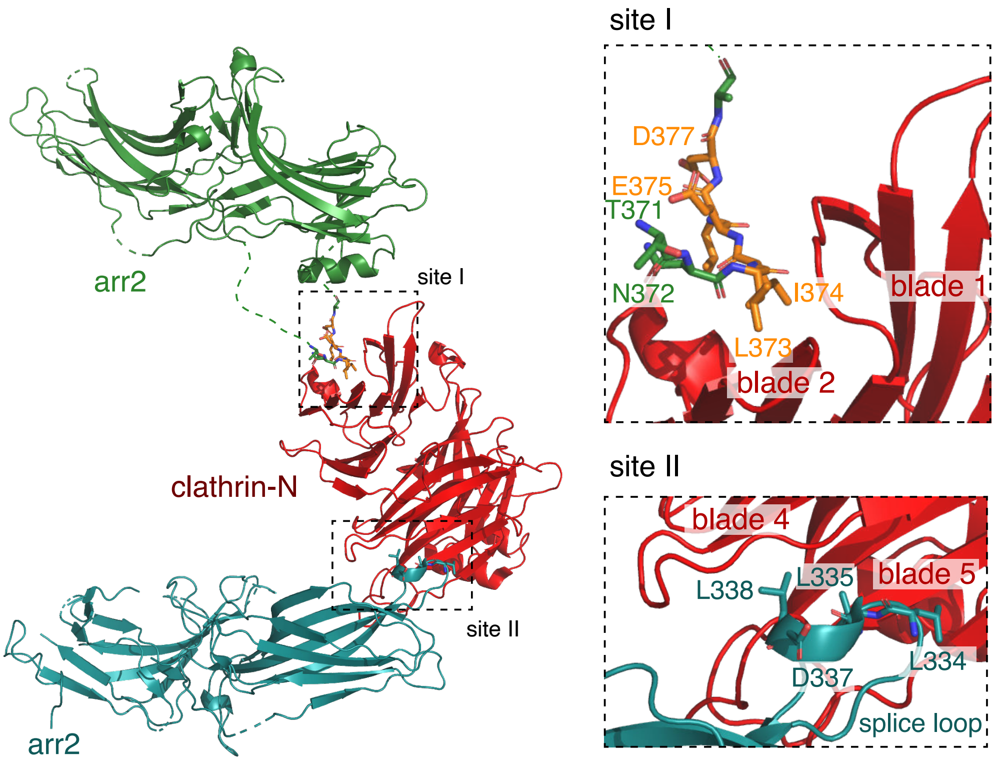


**Figure 1–figure supplement 1.** Arrestin sequence alignment and analysis of the arrestin2-clathrin-N complex structure. (A) Sequence alignment of four human arrestin types. The clathrin-binding motif (orange) and AP2 binding motif (purple) are indicated within the sequences. The numbering on the right corresponds to the last amino acid in the alignment. (B) X-ray structure of the arrestin2-clathrin complex (PDB:3GD1[45]). The structure shows two arrestin chains (green and blue) interacting with one clathrin-N molecule (red). Interaction sites 1 and 2 are indicated by dashed rectangles with enlarged views on the right. The arrestin2 residues involved in the interaction are shown as sticks. Site I: between CBL of arrestin2 (green) and the edge of β-sheet blade 1 and 2 of clathrin-N. CBL residues are shown as sticks. The numbering follows the PDB deposition. Site II: between arrestin2 splice loop (blue) and β-sheet blade 4 and 5 of clathrin-N.

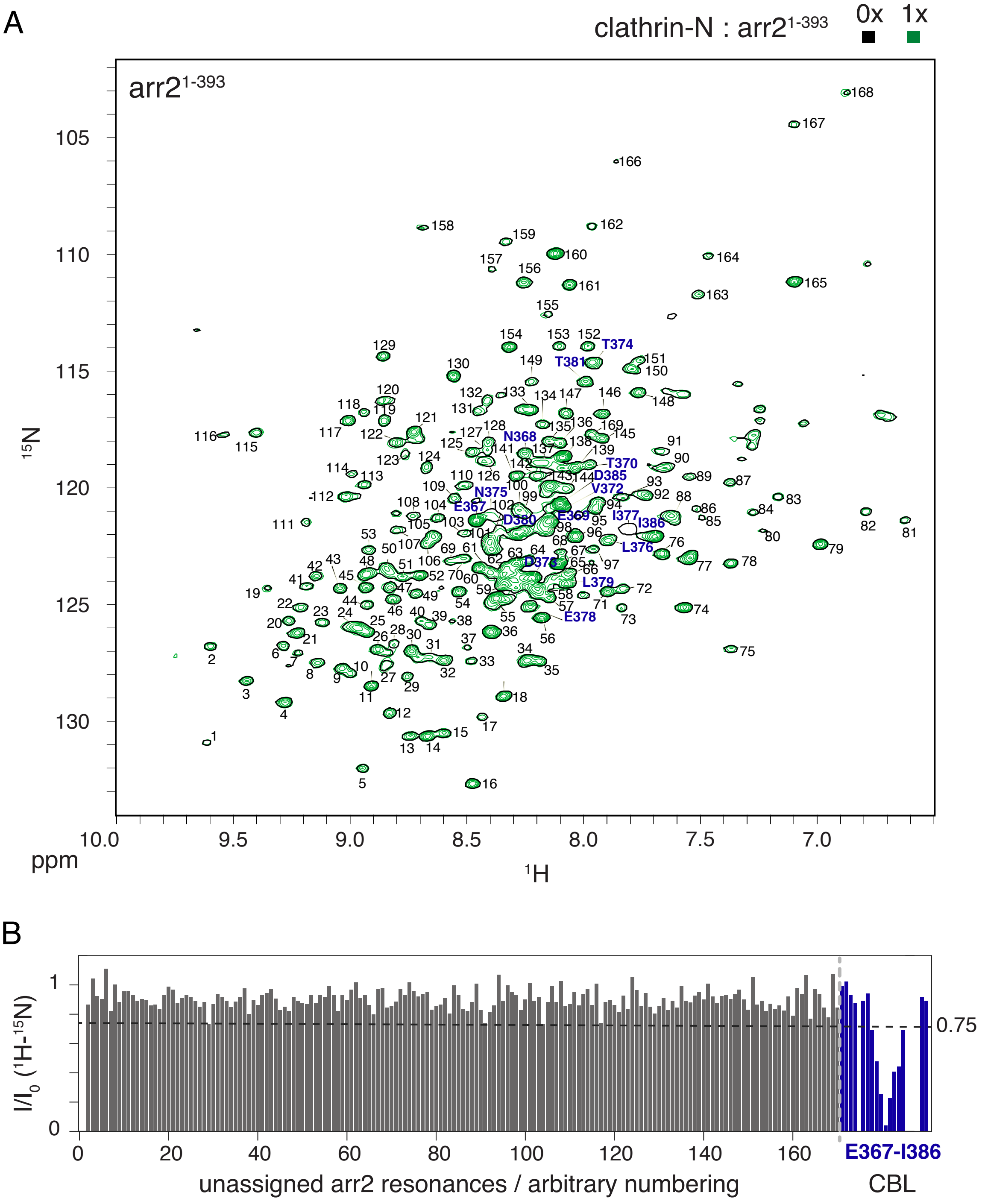


**Figure 1–figure supplement 2.** Mapping of clathrin-N interaction on arrestin2. (A) $^{1}H$-$^{15}N$ TROSY spectrum of arrestin2$^{1-393}$ in apo (black) or upon equimolar addition of clathrin-N (green) recorded on a Bruker AVANCE 14.1 T (600 MHz) spectrometer equipped with a TCI cryoprobe at 303 K. Assigned resonances are marked by blue amino acid numbers. Unassigned resonances are marked by black arbitrary numbers. (B) Intensity reduction $I/I_0$ of arrestin2$^{1-393}$ $^{1}H$-$^{15}N$ TROSY resonances upon clathrin-N interaction. The dashed line is drawn at one standard deviation below the average $I/I_0$ value. The resonance numbering follows panel (A).

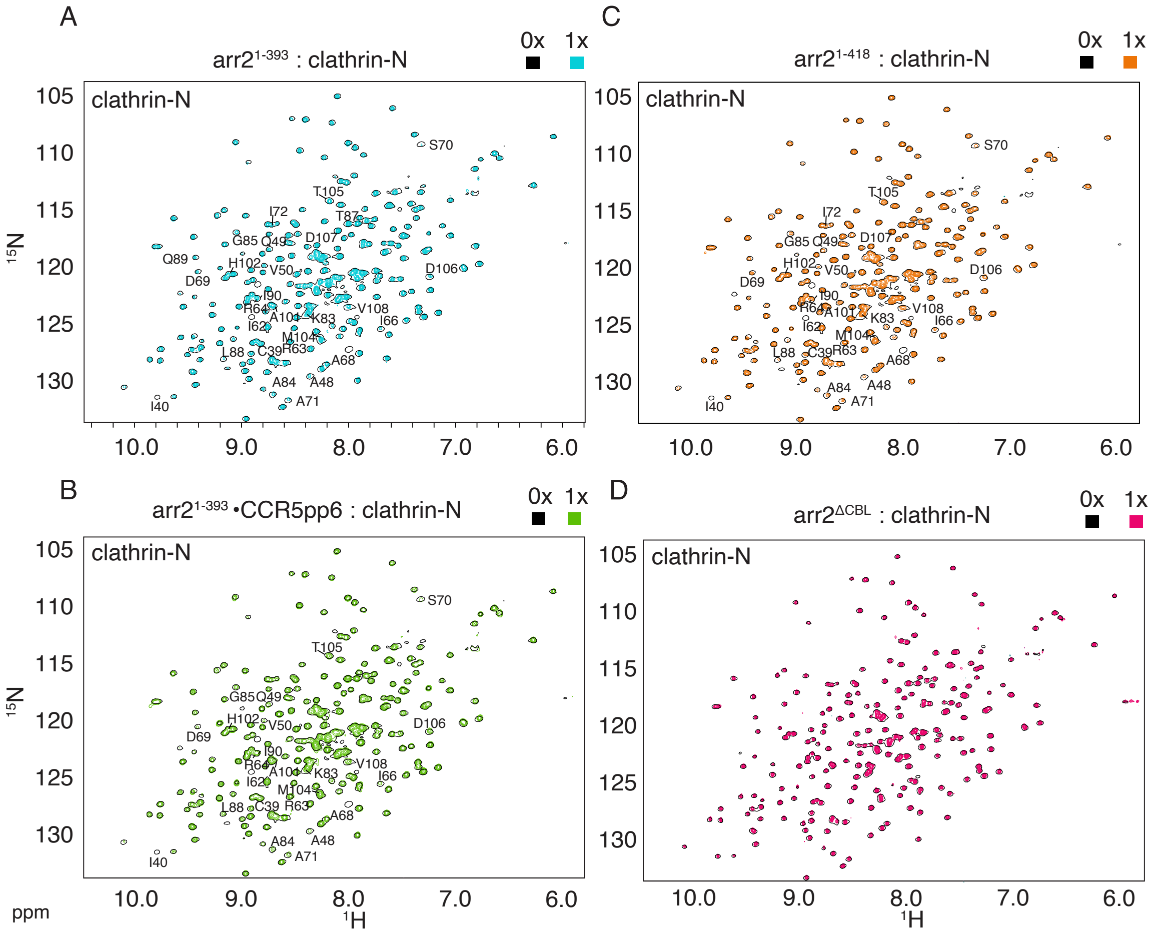


**Figure 2–figure supplement 1.** Mapping of arrestin2 interaction on clathrin-N. $^{1}$H-$^{15}$N TROSY spectra of clathrin-N in the presence of one molar equivalent of (A) arrestin2$^{1\text{-}393}$ (cyan); (B) arrestin2$^{1\text{-}393}$•CCR5pp6 (green); (C) arrestin2$^{1\text{-}418}$ (orange) or (D), arrestin2$^{\Delta CBL}$ (red) superimposed on $^{1}$H-$^{15}$N TROSY spectrum of apo clathrin-N (black). The spectra were recorded on a Bruker AVANCE 21.1 T (900 MHz) spectrometer equipped with a TCI cryoprobe at 303 K. Assigned residues undergoing significant line broadening are indicated on the spectra.

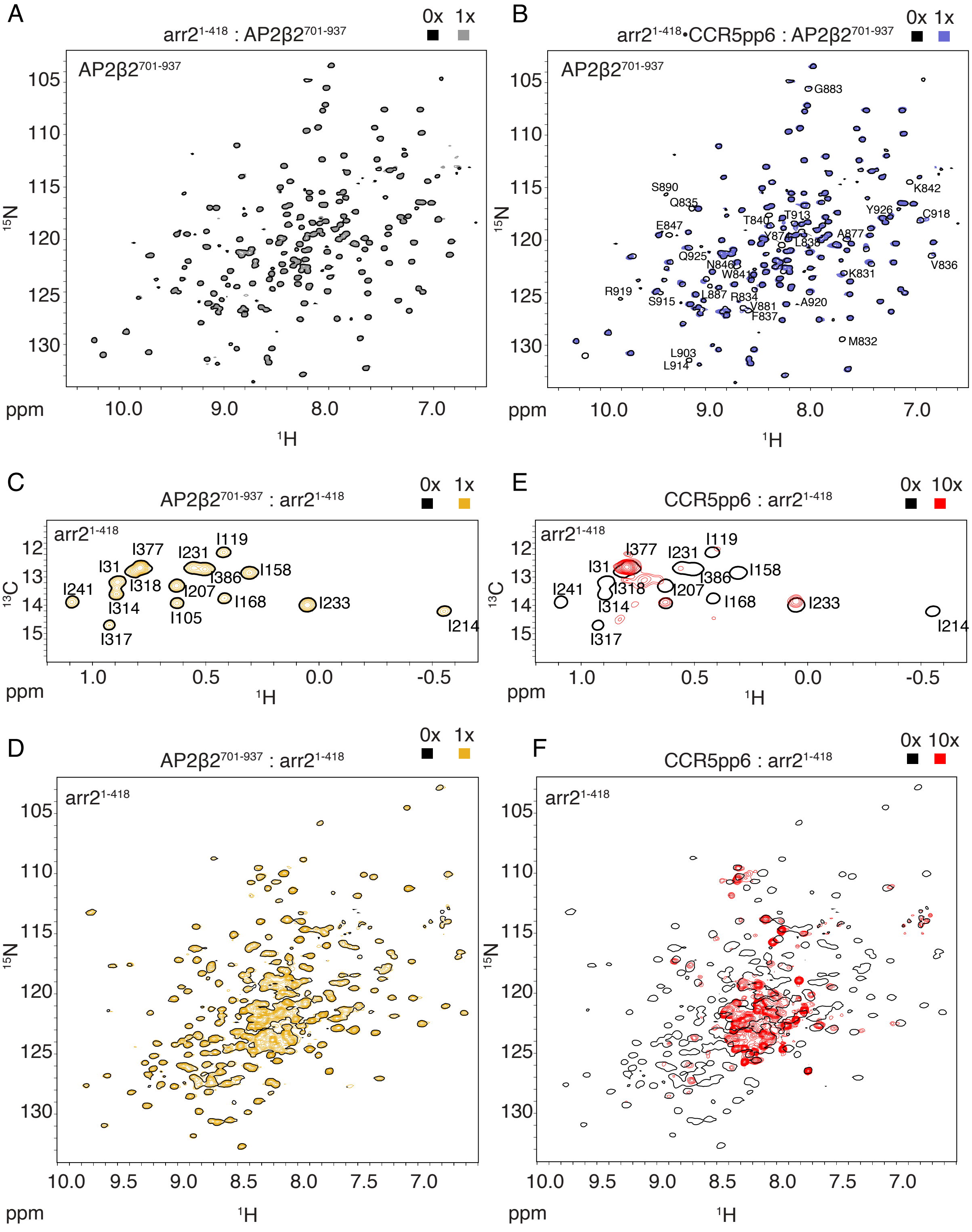


**Figure 3 – figure supplement 1.** Arrestin2 interaction with the C-terminal domain of AP2β2 detected by NMR. (A, B) $^{1}$H-$^{15}$N TROSY spectra of AP2β2$^{701\text{-}937}$ in the absence or presence of one molar equivalent of either arrestin2$^{1\text{-}418}$ (A) or arrestin2$^{1\text{-}418}$•CCR5pp6 (B). Assigned residues undergoing significant intensity reduction are indicated on the spectra. (C, D) Test of interaction of arrestin2$^{1\text{-}418}$ with AP2β2$^{701\text{-}937}$. (C) $^{1}$H-$^{13}$C HMQC (Ile-δ1-$^{13}CH_3$, $^{2}$H-labeled arrestin2$^{1-418}$) and (D) $^{1}H$-$^{15}N$ TROSY ($^{15}N$, $^{2}H$-labeled arrestin2$^{1-418}$) spectra of arrestin2$^{1-418}$ in the absence or presence of one molar equivalent of AP2β2$^{701-937}$. (E, F) Test of interaction of arrestin2$^{1-418}$ with CCR5pp6. (E) $^{1}H$-$^{13}C$ HMQC (Ile-δ1-$^{13}CH_3$, $^{2}H$-labeled arrestin2$^{1-418}$) and (F) $^{1}H$-$^{15}N$ TROSY ($^{15}N$, $^{2}H$-labeled arrestin2$^{1-418}$) spectra of arrestin2$^{1-418}$ in the absence or presence of a ten-molar equivalent of CCR5pp6.

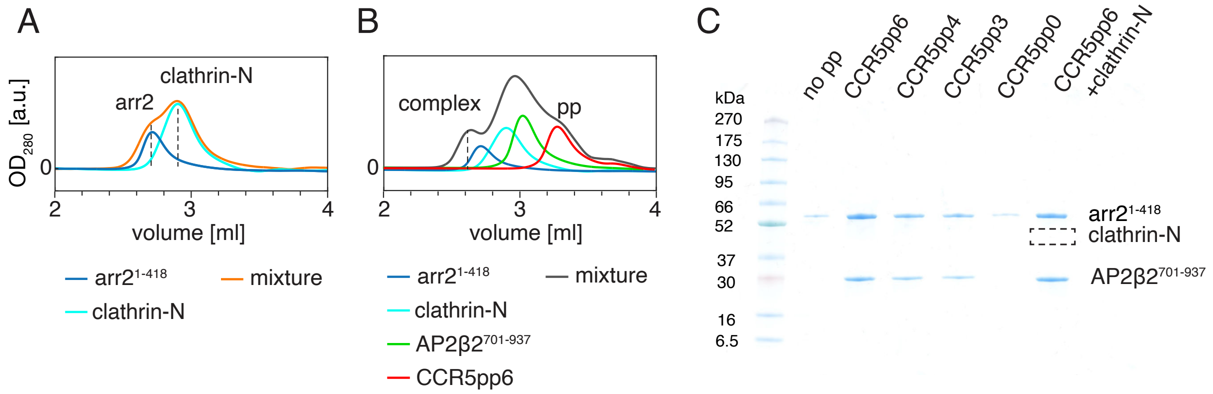


**Figure 3–figure supplement 2.** Arrestin2 interaction with clathrin-N and AP2β2 monitored by SEC. (A) SEC profiles of arrestin2$^{1\text{-}418}$, clathrin-N and their mixture. (B) as in (A) but in the presence of AP2β2$^{701\text{-}937}$ and CCR5pp6. (C) SDS-PAGE of sample from the SEC complex fraction (taken at 2.6ml) monitoring the formation of the arrestin2/AP2β2 complex for various conditions indicated on the top. The empty square indicates the expected position of clathrin-N.

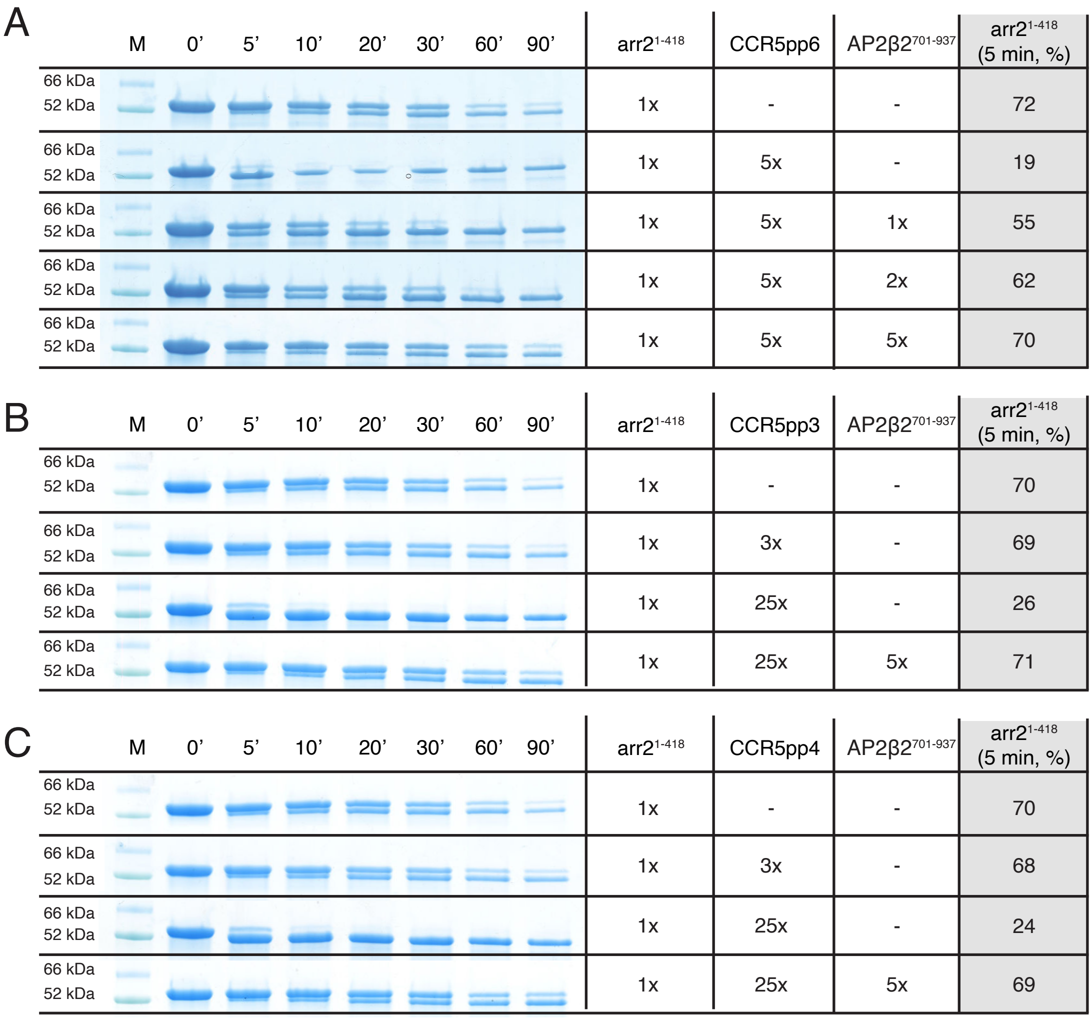


**Figure 3–figure supplement 3.** Arrestin2$^{1\text{-}418}$ trypsin proteolysis assay. Trypsin proteolysis of arrestin2$^{1\text{-}418}$ in apo form and in complexes with (A) CCR5pp6, (B) CCR5pp3 and (C) CCR5pp4 and AP2β2$^{701\text{-}937}$ detected by SDS-PAGE. The incubation time is indicated on the top. ‘M’ indicates the protein size marker. The table on the right indicates sample composition and the relative abundance of arrestin2$^{1\text{-}418}$ after 5 min. incubation (see Materials and Methods). The activation by the CCR5 phosphopeptides accelerates arrestin2$^{1\text{-}418}$ proteolysis. The addition of AP2β2$^{701\text{-}937}$ reverses this effect.

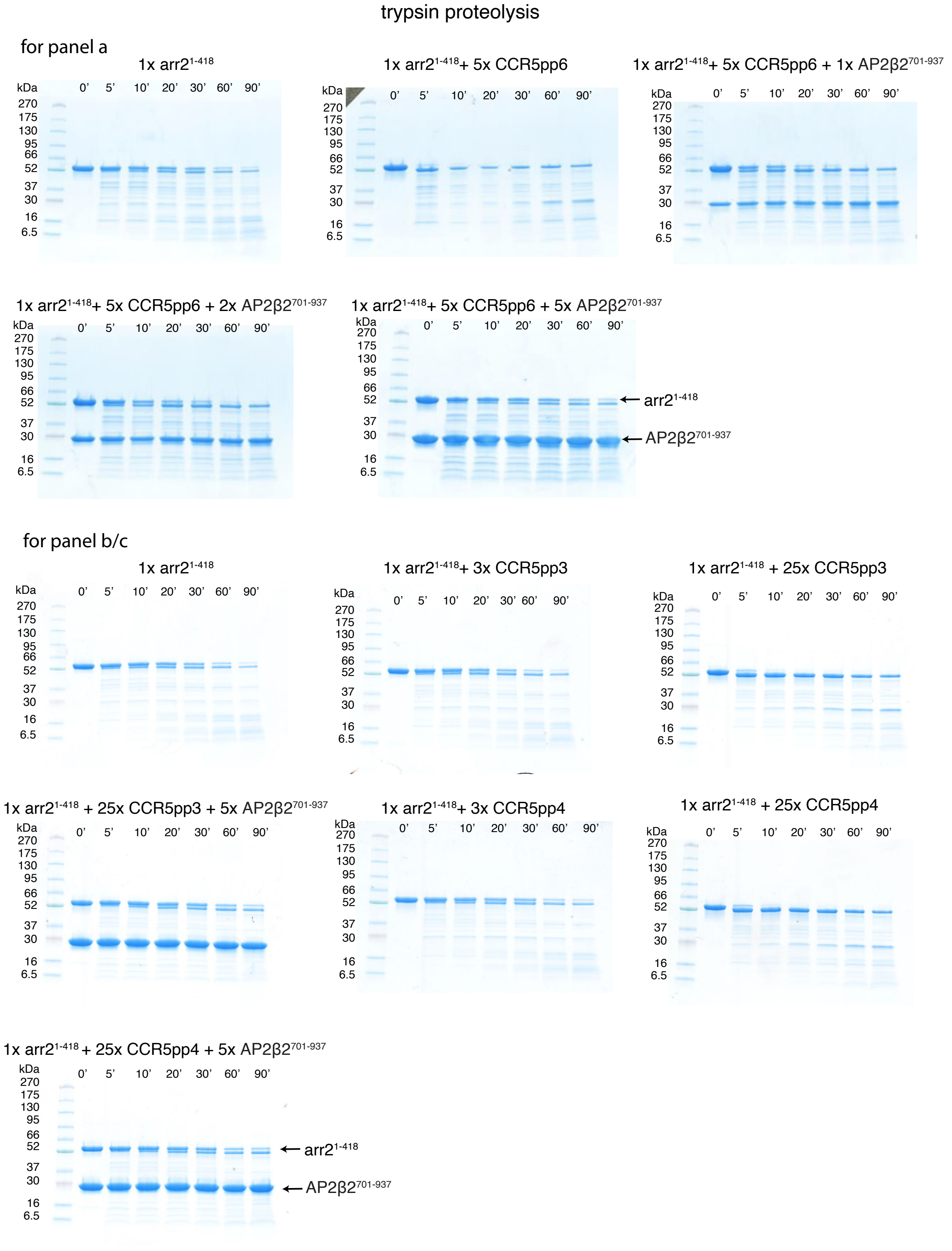


**Figure 3–figure supplement 4.** Uncropped SDS-PAGE gels of the arrestin2$^{1-418}$ trypsin proteolysis. SDS-PAGE gels for the data in Figure 3–figure supplement 3. The composition of the sample and the reaction incubation time are indicated at the top. Protein marker bands are indicated on the left. Arrestin2$^{1-418}$ runs at ~ 52 kDa, AP2β2$^{701-937}$ at ~ 30 kDa.

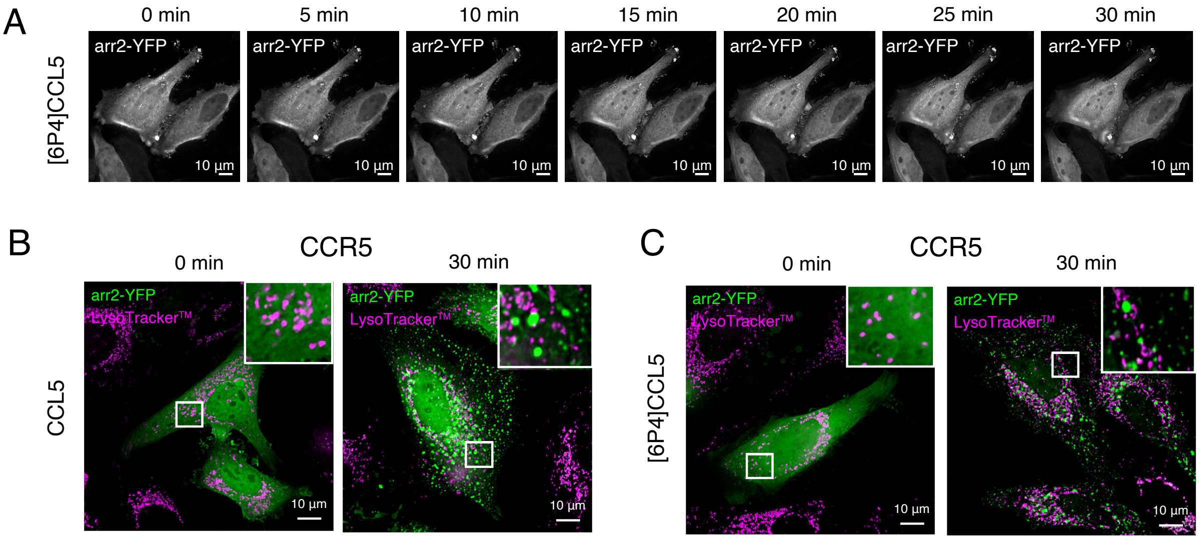


**Figure 4–figure supplement 1.** Arrestin2 internalization (control) and re-localization in live HeLa cells upon chemokine stimulation in the presence of CCR5. (A) HeLa cells transfected only with arrestin2-YFP and stimulated with 6P4[CCL5]. In the absence of the receptor, arrestin2 stays distributed in the cell cytoplasm and nucleus. (B,C) Arrestin2-YFP re-localization in HeLa cells transfected with wild-type CCR5 and arrestin2-YFP genes. Arrestin2-YFP and LysoTrackerTM signals were used to monitor the protein trafficking for up to 30 min after CCL5 (B) or 6P4[CCL5] (C) incubation. No overlap between the LysoTrackerTM and arrestin2 signal has been detected. White squares indicate zoomed regions of interest.

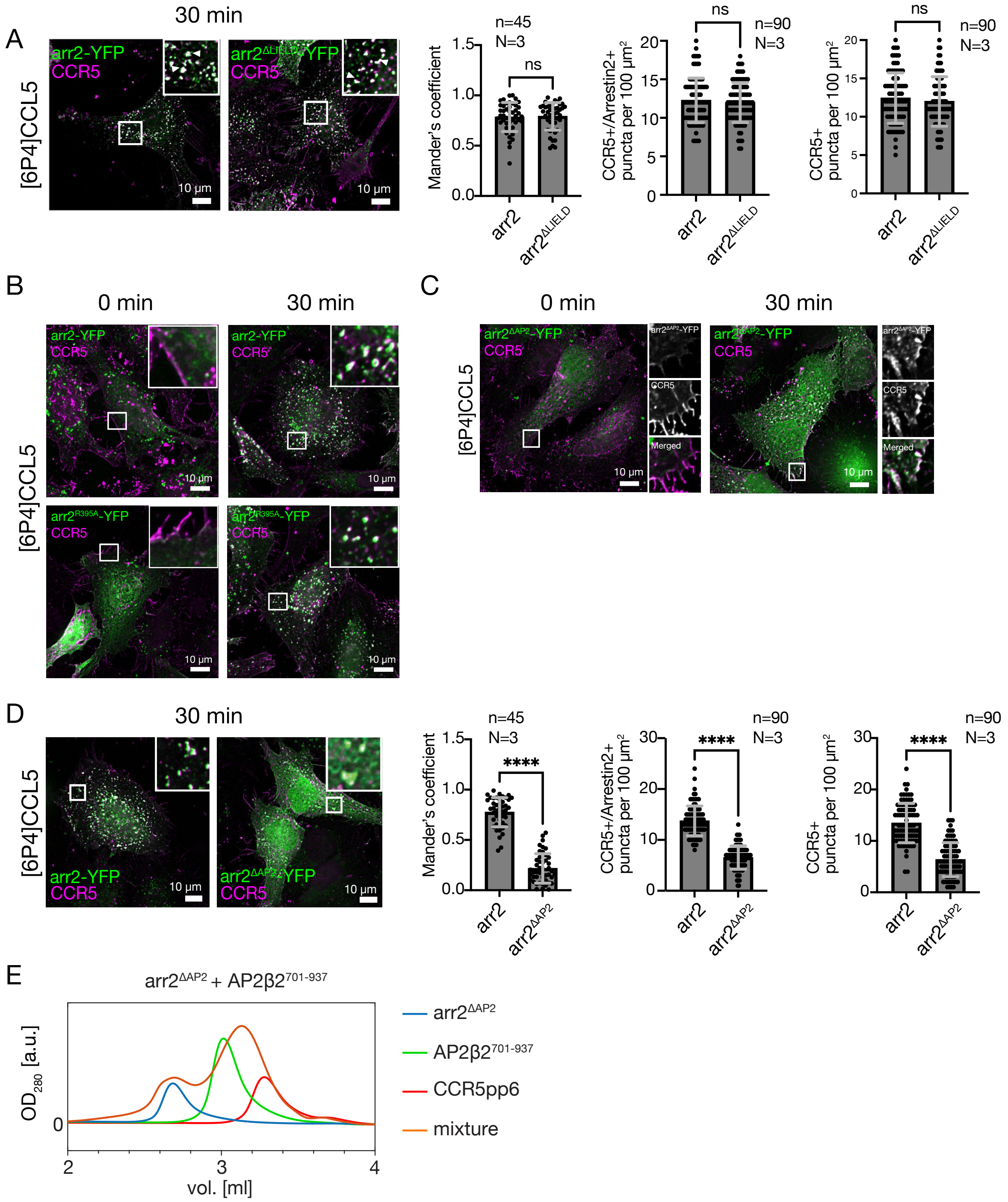


**Figure 5–figure supplement 1.** Dependence of CCR5 internalization on the interactions of arrestin2 with clathrin or AP2 monitored in HeLa (CCL2) cells by immunofluorescence microscopy. (A) CCR5 internalization induced by [6P4]CCL5 in HeLa cells co-transfected with plasmids containing either arrestin2-YFP or arrestin2$^{\Delta LIELD}$-YFP, which misses the clathrin-binding motif. No significant difference is detected as quantified by the Mander's colocalization coefficient (ns, not significant: P > 0.9999) or by counting CCR5$^{+}$/Arrestin2$^{+}$ and only CCR5$^{+}$ puncta 30 min after ligand incubation. Mean and standard deviation are shown for N=3

biological replicates and n=45 ROI from 20 cells (Mander's) or n=90 ROI from 30 cells (puncta counting). (B) CCR5 internalization induced by 6P4[CCL5] in HeLa cells co-transfected with plasmids containing either arrestin2-YFP or arrestin2$^{R395A}$-YFP, which bears a single mutation in the adaptin-binding motif. The mutation does not affect CCR5 internalization up to 30 min after 6P4[CCL5] stimulation. (C) Recruitment of arrestin2 $^{\Delta AP2}$-YFP to CCR5 in the HeLa cell plasma membrane upon [6P4]CCL5 ligand stimulation. (D) CCR5 internalization induced by [6P4]CCL5 in HeLa cells co-transfected with plasmids containing either arrestin2-YFP or arrestin2$^{\Delta AP2}$-YFP. Mean and standard deviation of the Mander's coefficient and the number of puncta were determined as in (A). The absence of the AP2 binding motif in the arrestin2$^{\Delta AP2}$-YFP construct causes a significant (****: $P < 0.0001$) reduction of CCR5 internalization 30 min after ligand incubation. (E) SEC profiles of arrestin2$^{\Delta AP2}$, AP2β2$^{701\text{-}937}$, CCR5pp6 and their mixture. No complex formation is observed.
